# *Chlamydia* pan-genomic analysis reveals balance between host adaptation and selective pressure to genome reduction

**DOI:** 10.1101/506121

**Authors:** Olga Sigalova, Andrei V. Chaplin, Olga O. Bochkareva, Pavel V. Shelyakin, Vsevolod A. Filaretov, Evgeny E. Akkuratov, Valentina Burskaya, Mikhail S. Gelfand

## Abstract

*Chlamydia* are ancient intracellular pathogens with reduced, though strikingly conserved genome. Despite their parasitic lifestyle and isolated intracellular environment, these bacteria managed to avoid accumulation of deleterious mutations leading to subsequent genome degradation characteristic for many parasitic bacteria.

We report pan-genomic analysis of eleven species from genus *Chlamydia* including identification and functional annotation of orthologous genes, and characterization of gene gains, losses, and rearrangements. We demonstrate the overall genome stability of these bacteria as indicated by a large fraction of common genes with conserved genomic locations. On the other hand, extreme evolvability is confined to several paralogous gene families such as polymorphic membrane proteins and phospholipase D and likely is caused by the pressure from the host immune system.

This combination of a large, conserved core genome and a small, evolvable periphery likely reflect the balance between the selective pressure towards genome reduction and the need to adapt to changing host environment.

## Introduction

Bacteria of genus *Chlamydia* are intracellular pathogens of high medical significance. They are important agents of sexually transmitted disease and pneumonia worldwide as well as the main cause of preventable blindness in developing countries (Elwell, Mirrashidi, and Engel, 2016; Nunes and Gomes, 2014). At that, no vaccines exist against human chlamydial strains, the recurrence rate is high, and persistent chlamydial infections are associated with a higher risk of atherosclerosis, reactive arthritis, and oncogenic effects (Campbell and Rosenfeld, 2014; Chumduri et al., 2013; Villareal, Whittum-Hudson, and Hudson, 2002; ECDC, 2015). The number of cases does not decrease with better examination schemas. In addition, the genus includes zoonotic pathogens which cause life-threatening diseases if infecting humans (Nunes and Gomes, 2014; ECDC, 2015).

These bacteria have a complex biphasic lifecycle (AbdelRahman and Belland, 2005), which implies changes in the DNA compaction (Barry, Hayes, and Hackstadt, 1992), metabolism (Omsland et al., 2014), and temporal expression of early, middle, and late genes (Shaw et al., 2000). Extracellular forms of *Chlamydia* (*elementary bodies*) attach to host cells and initiate endocytosis, yielding formation of a membrane-bound compartment, the *inclusion body*. Once inside the inclusion, elementary bodies differentiate into a larger intracellular form (*reticulate bodies*), engaging into complex host-pathogen interactions. In particular, these bacteria reorganize vesicular transport, prevent apoptosis, slow down the host cell cycle, and suppress inflammatory immune response by damping nuclear factor B transcription, for a review see (Elwell, Mirrashidi, and Engel, 2016). The chlamydial inclusion membrane has been referred to as a pathogen-specified parasitic organelle (Moore and Ouellette, 2014).

As summarized in (Koonin and Wolf, 2008), a typical bacterial genome is shaped by the dynamic interaction of six major evolutionary forces directed towards either genome reduction or complexification. Genome streamlining and degradation lead to genome contraction under strong purifying or neutral selection, respectively. This is counteracted by genome complexification and innovation via gene duplications, operon shuffling, horizontal gene transfer (HGT), or propagation of mobile elements. And although all these forces might act simultaneously, their contribution to shaping individual prokaryotic genomes is strikingly different, reflecting the ecological niche and the lifestyle of a microorganism.

As a consequence of their obligatory intracellular lifestyle, *Chlamydia* have reduced genomes of about 1 Mb and 850–1100 genes. At most 14 transcription factors (TFs) have been predicted to regulate gene expression (Domman, and Horn, 2015). However, as opposed to many other pathogens with reduced genomes (Moran, 2002), this apparent simplification likely resulted from genome streamlining rather than degradation. In particular, genomes of all chlamydial species have a low number of pseudogenes (Nunes and Gomes, 2014; Bachmann, Polkinghorne, and Timms, 2014). The gene order is highly conserved everywhere outside the *plasticity zone*, a genomic region of about 81 kB around the replication terminus (Read et al., 2000). Similarly, the gene content is conserved, with the majority of genes being shared with other representatives of the phylum (Collingro et al., 2011). Finally, genomes of *Chlamydia* spp. are generally free of disruptive mobile elements, with the exception of the IS-associated tetracycline-resistance genomic island in *C. suis* and the remnants of IS-like elements and prophages in some genomes (Vorimore et al., 2013; Sachse et al., 2014; Sachse et al., 2015; Nunes and Gomes, 2014).

On the other hand, the genome of *Chlamydia* is evolvable enough to allow for significant phenotypic variation with twelve currently recognized species having a broad range of host specificities and tissue tropisms (Sachse et al., 2015; Vorimore et al., 2013; Sachse et al., 2014). In particular, *C. trachomatis* are exclusively human pathogens causing trachoma (serovars A-C), lymphogranuloma venereum (LGV, serovars L1-L3), and epithelial urogenital infections (serovars D-K). *C. pneumoniae* is one of the most common causes of respiratory infections in humans, also able to infect animals, e.g. horses, marsupials, and frogs. Other species infect a broad range of animals including mice, guinea pigs, birds, cattle, sheep, swine, horses, cats, koalas, frogs, and snakes, causing various diseases, which in some cases can be transmitted to humans, for a review see (Nunes and Gomes, 2014). The source of this phenotypic diversity is limited, and most differences in the host specificity and tissue tropism have been attributed to several highly variable gene families including cytotoxin (Belland et al., 2001), polymorphic outer membrane proteins (Gomes et al., 2006), inclusion membrane proteins (Dehoux et al., 2011), and phospholipase D enzymes (Nelson et al., 2006), as well as several metabolic pathways, such as tryptophan (Caldwell et al., 2003) and biotin (Nunes and Gomes, 2014) biosynthesis.

The pan-genome approach (Tettelin et al., 2005) is a comparative genomic technique which implies analysis of conservation and evolution of individual gene families by pooling together complete genomes from a clade of microorganisms. The pan-genomic analysis implies that all genes from a group of closely related prokaryotic genomes are pooled together and then clustered into orthologous groups (OGs) by sequence similarity. OGs are then classified into universally conserved (“core”), non-universal (“periphery”), and unique (“cloud” or “singletons”) gene fractions. The structure of the pan-genome provides a robust description of the phylogenetic structure within a prokaryotic group (Gordienko, Kazanov, and Gelfand, 2013; Moldovan and Gelfand, 2018) as well as the lifestyle of microorganisms from the group (Rouli et al., 2015). Due to the high medical relevance of *Chlamydia*, the research on their genomes has greatly expanded over the last decade. Previous studies include pan-genomic analysis of phylum *Chlamydiae* (Collingro et al., 2011) and order *Chlamydiales* (Psomopoulos et al., 2012), as well as multiple comparative studies of individual chlamydial species or gene families.

In this study, we provide a comprehensive pangenomic analysis of 127 strains from twelve species of genus *Chlamydia* and assess the contribution of genome reduction and complexification processes in shaping chlamydial genome. The paper is organized as follows. First, we identify and functionally characterize the conserved and variable components of the chlamydial pan-genome. Next, we focus on the interplay between the genome complexification and streamlining via the analysis of gene losses, pseudogenes, genomic rear-rangements, and expansion of paralogous gene families. Finally, we provide a case-study of the Polymorphic membrane protein G (PmpG) gene family representing an interesting combination of extensive paralogisation, phase and antigen variation, and pseudogenisation.

## Results

### Pan-genome of genus *Chlamydia*: large core, small periphery, and tiny cloud

The pan-genome of 127 genomes from twelve species of *Chlamydia* consisted of 725 core, 1185 peripheral, and 685 singleton orthologous groups (Table 1). The set of sequenced genomes in the NCBI database was highly biased with the majority of strains coming from *trachomatis*, followed by *C. psittaci* and *C. pneumoniae* reflecting the high medical significance of these species. To check whether the pan-genome definition is influenced by the sample biases, we reproduced the same analysis limiting the sample size to one or two strains per species. This resulted in about 8% of OGs (mostly from paralogous gene families) changing their composition, which was compatible with the results obtained upon varying the default orthoMCL parameters. Therefore, we conclude that the pan-genome provides a robust representation of the chlamydial gene repertoire. An additional round of clustering was performed to reveal relationships between different orthologous groups. As a result, 508 orthologous groups were grouped into 157 clusters (referred to as *orthologous families*, or OFs) with 27 OFs containing four or more OGs (Table 1). In particular, 156 singletons clustered into OFs with other genes from the core or the periphery, suggesting their origin via paralogization and/or functional divergence from the common pool of genes.

**Table 1:** Summary of the pan-genome properties.

|  | Number of OGs | with length exceeding 100 aa | ...with Pfam annotation | ...with GO annotation |
| --- | --- | --- | --- | --- |
| Core genome | 725 | 692 | 629 | 443 |
| Periphery | 1185 | 454 | 245 | 100 |
| Singletons | 685 | 76 | 36 | 10 |

In agreement with previous studies, our analysis shows that the gene repertoire of *Chlamydia* is highly optimised in that it is comprised of a large core, small periphery, and tiny cloud. Firstly, out of about 1000 protein-coding genes in an average chlamydial genome, 75% were universally conserved through the whole genus. The median length of the encoded proteins is 307 amino acids which is typical for prokaryotic genomes (Xu et al., 2006), and most of them have conserved domains according to the Pfam database (Table 1). Secondly, the chlamydial periphery is mostly species-specific with low overall genomic diversity within species (Figure 1). Two peaks at 81 and 46 genomes represent orthologous groups specific to two species groups previously assigned to genera *Chlamydia* and *Chlamydophila*, a division proposed by (Everett, Bush, and Andersen, 1999) but not generally accepted by the research community (Stephens et al., 2009). An additional small plateau approximately at 20 strains is formed by genes unique to the monoplyletic clade comprised of *C. psittaci*, *C. abortus*, *C. felis*, and *C. caviae*.

**Figure 1:**
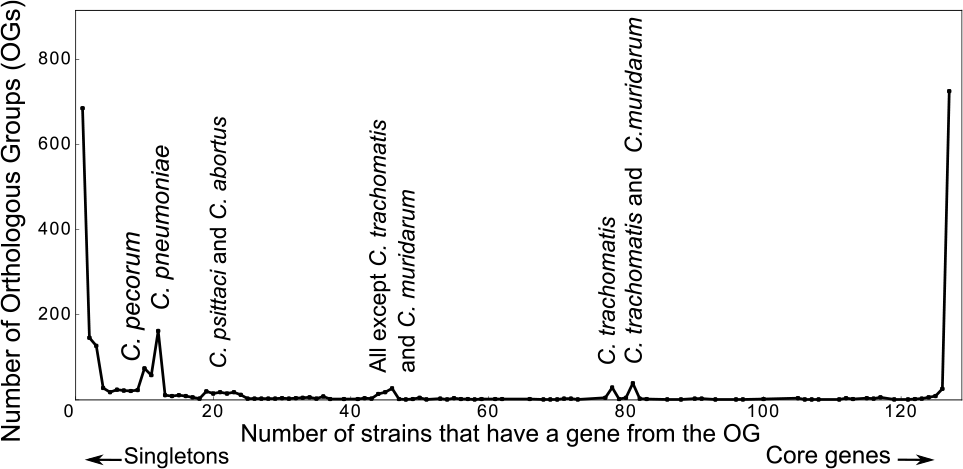
Distribution of Orthologous Groups (OGs) by their presence in different subsets of Chlamydia strains (the U-curve) The left and right peaks of the curve correspond to unique (singletons) and universally conserved (core) genes, respectively. The internal peaks are mainly formed by OGs specific to individual species or monophyletic groups of species. Pan-genomes of individual species yield plots similar to that of the genus pan-genome, but without intermediate peaks (data not shown).

Finally, the majority of singletons and about half of the periphery genes are short reading frames, less than 100 amino acids long and without functional annotation, and hence may result from overprediction by gene-calling algorithms. Even without accounting for this possibility, the cloud of chlamydial pan-genome is tiny with just about 5 unique genes per genome.

The pan-genome of genus *Chlamydia* is open, meaning that we expect the overall pan-genome size to increase with the addition of new genome sequences (Figure 2). Indeed the number of new genes decreases with each new genome *n* at the rate *N* (*n*) = 215*n*^0,82^confirming that the pan-genome is indeed open (Supplementary file Figure 1A) and the Chao lower bound estimate (Chao, 1987) of the pan-genome size is 4189 genes. However, the percentile pan-genomes (Gordienko, Kazanov, and Gelfand, 2013) corresponding to OGs present in at least a given fractions of strains, saturate after just a few initial strains (Figure 2B). This shows that the pan-genome growth is mainly due to singletons and rare OGs. Further exclusion of short reading frames (less than 300 bp) leads to a marked decrease of the pan-genome growth rate (Figure 2C). The rate of gene gain with new genomes becomes *N*(*n*) = 136*n*^1,14^ (Supplementary file Figure 1B), what corresponds to a closed pan-genome, and the Chao estimate decreases to 1397 genes. Clustering of the remaining OGs into orthologous families makes the pan-genome of the whole genus closed despite high heterogeneity of the sample (Figure 2D).

**Figure 2:**
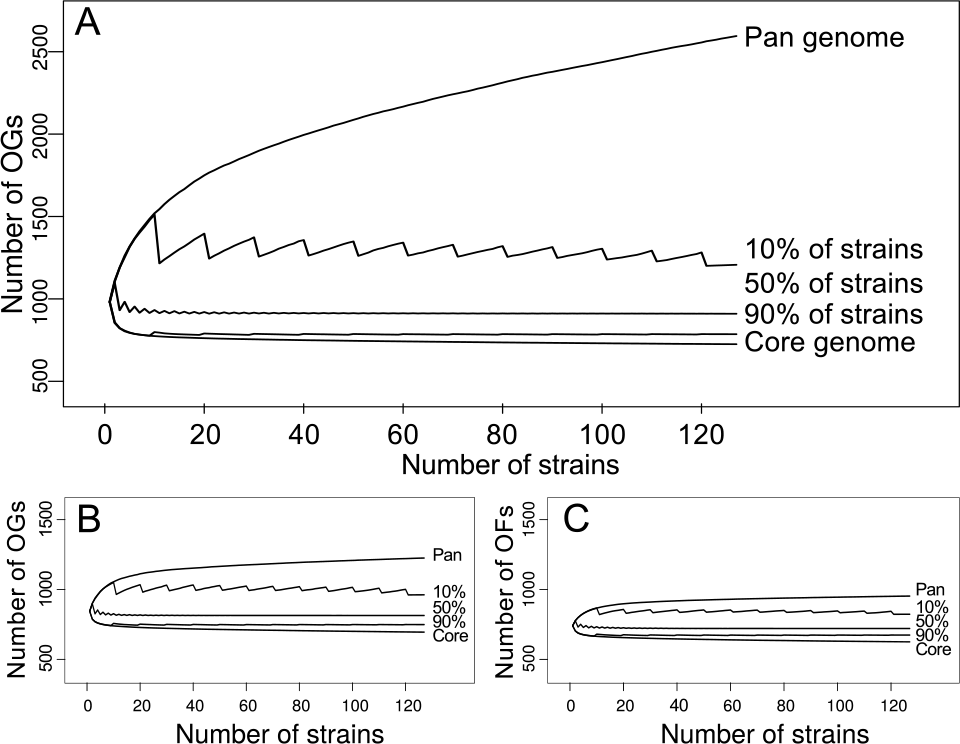
Numbers of Orthologous Groups (OGs) present in a given fraction of chlamydial genomes as dependent on the number of sequentially added genomes (saturation curves) The topmost curve shows the pan-genome size, the lowest curve shows the core genome size, and the remaining curves show the percentile pan-genome sizes. Each dot represents a mean value obtained from 500 random permutations of the strain addition order. The jagged pattern is a consequence of the rounding procedure. Panels:• Plot for all OGs of Chlamydia spp. (B) Plot for OGs with the median length exceeding 100 amino acids. (C) Plot for all OGs with the median length exceeding 100 amino acids and clustered into Orthologous Families.

### Functional annotation shows low diversity of the *Chlamydia* metabolism

The chlamydial core genome is mainly comprised of the machinery for processing the genetic information and enzymes of the central metabolism while the periphery is enriched in virulence-related gene families such as inclusion membrane proteins and polymorphic membrane proteins (Figure 3).

**Figure 3:**
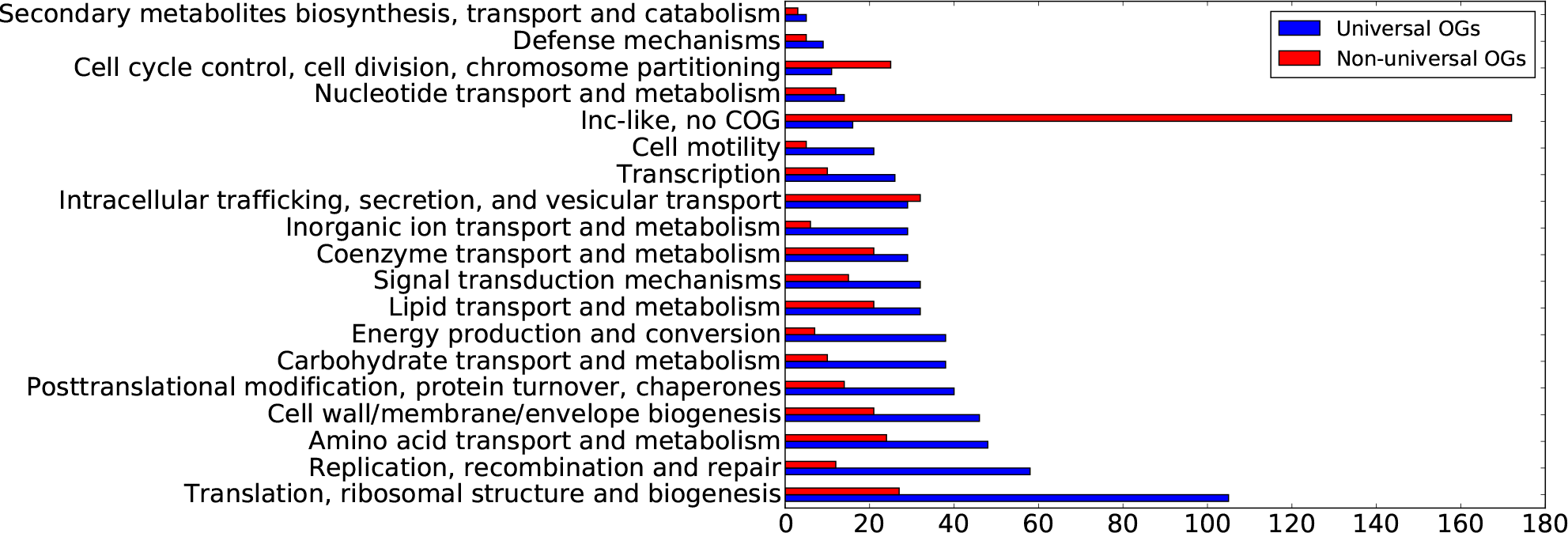
Functional annotation of orthologous groups. Clusters of Orthologous Genes (COGs) assigned to ten or less orthologous groups are not shown.

All studied strains lack pathways to synthesize the majority of amino acids while presumably preserving mechanisms for performing glycine-serine and glutamate-aspartate interconversions (encoded by serine hydroxymethyltransferase and putative aspartate aminotransferase genes, respectively). The energy metabolism is similar among all *Chlamydia*. All studied strains have a complete glycolysis pathway with the exception of hexokinase, and also the hexose phosphate transporter HPTcp, thus allowing *Chlamydia* to use glucose-6-phosphate as an energy source (Omsland et al., 2012; Schwöppe, Winkler, and Neuhaus, 2002). ADP/ATP Translocase NTT1, providing an alternative pathway to obtain ATP (Tjaden et al., 1999), is present in all *Chlamydia*, and only in one genome (*Chlamydia psittaci* 84/55) the corresponding gene has a single nucleotide insertion in its coding sequence presumably leading to production of a nonfunctional truncated product. All studied *Chlamydia* strains harbour three genes encoding heat-shock protein GroEL (McNally and Fares, 2007; Karunakaran et al., 2003).

Some very common protein families are absent in the chlamydial pan-genome. For example, all *Chlamydia* lack GTP diphosphokinases (Pfam domain families PF04607 and PF13328) and are probably unable to regulate gene expression by (p)ppGpp. Moreover, *Chlamydia* have also lost the omega subunit of RNA polymerase (PF01192) known to participate in binding (p)ppGpp (Ross et al., 2013; Syal and Chatterji, 2015). All *Chlamydia* lack cell division protein FtsZ (PF12327, PF00091) and classical peptidoglycan transglycosylase domains (PF00912), which is not surprising in the light of the unique division mechanism of *Chlamydia* (Jacquier, Viollier, and Greub, 2015; Liechti et al., 2016). Another notable loss, which could be explained by stable conditions within host cells, is cold shock proteins (PF00313) that weakly bind to single stranded RNA and destabilize RNA secondary structures (Goldstein, Pollitt, and Inouye, 1990; Doniger et al., 1992).

Non-universal genes were mapped to several metabolic pathways, including biosynthesis of biotin, tryptophan, thiamine, folate, purines, and pyrimidines. In particular, *C. trachomatis*, *C. pecorum*, and *C. muridarum* strains lack phenylalanine hydroxylase responsible for the biosynthesis of tyrosine from phenylalanine. *C. pecorum*, *C. felis* and *C. caviae* have a complete pathway of tryptophan production from kynurenine, while genital strains of *C. trachomatis* are able to produce tryptophan from exogenous indole (Caldwell et al., 2003). Generally, the non-universal distribution of some enzymes from these pathways has been shown to play a role in tissue tropism and persistent infections (Read et al., 2003; Nunes and Gomes, 2014; Sait et al., 2014).

### Gene loss and genomic rearrangements did not contribute much to the Chlamydial phenotypic diversity

The number of species-specific gene losses is markedly low (Supplementary file Table 1) indicating that a major genome reduction has been completed prior to speciation, with a few exceptions like the loss of folic acid biosynthesis in *C. pecorum* (Sait et al., 2014). Similarly, there are very few cases of genes absent in any monophyletic clade in the species phylogenetic tree that could be explained by single gene loss (22 OGs if excluding former *Chlamydia/Chlamidophyla* split), while 182 OGs were monophyletically gained and 175 OGs have mosaic phyletic patterns (Supplementary file Figure 2). The latter could result from gene inflow by horizontal gene transfer (HGT), independent paralogisation events, or multiple parallel gene losses.

To discriminate gene inflow by HGT from the two other types of events, we applied the following model. If a gene with a mosaic phyletic pattern has been inherited vertically from the common ancestor and lost by several genomes, we expect to find it at the same syntenic region in the remaining strains. Genes not satisfying this condition are candidates for having been obtained horizontally. At that, only few cases of a mosaic location pattern for single-copied orthologous groups have been detected.

Eight genes related to the tryptophan biosynthesis and metabolism form an operon that has been found in the plasticity zone in *C. caviae* and *C. felis* and at a stable genome locus in *C. pecorum*; three of these genes are also found in *C. trachomatis*, while the genomes of other species lack all these genes (Figure 4).

**Figure 4:**
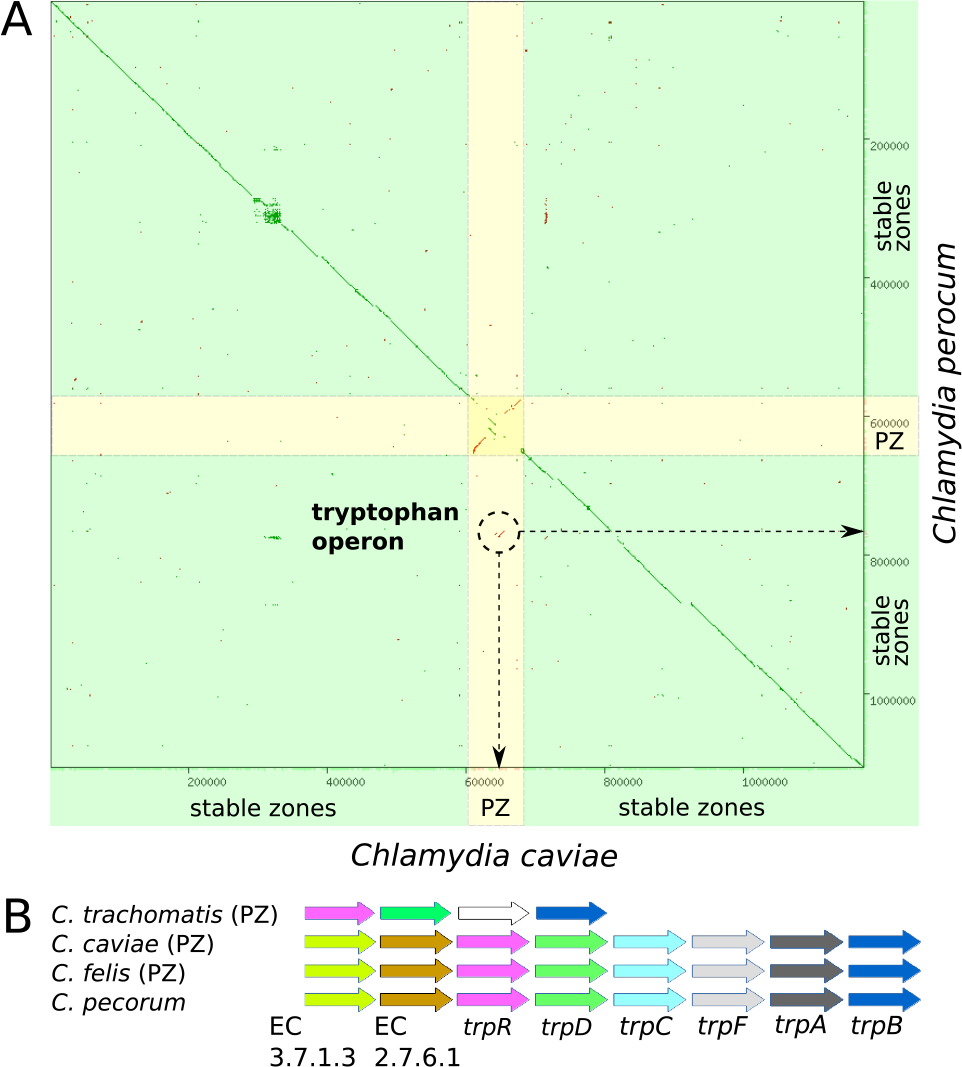
Dot-plot chromosome comparison of C. caviae and C. perocum revealing horizontal transfer of the DNA segment containing the tryptophan operon. PZ and yellow color denote the plasticity zone.

The only case likely involving multiple, independent gene transfers into different strains belonging to the same species is OG2167 encoding an outer membrane protein. The phylogenetic tree for this OG is inconsistent with the genus phylogeny, and this inconsistency is best explained by horizontal gene transfer between *C. psittaci* WS/RT/E30 and *C. abortus* LLG (Supplementary file Figure 3).

Therefore, the majority of polyphyletic OGs can not be explained by lateral gene transfer. Out of 175 OGs with a mosaic phyletic pattern, 82 are members of larger orthologous families (OFs) and hence their polyphyletic pattern can result from ancestral gene duplications and subsequent gene loss in some descendant strains. Remaining polyphyletic OGs with conserved genomic location are represented by single-copy genes having no paralogs within the genus and hence are most parsimoniously explained by parallel gene loss due to ongoing genome streamlining.

### Strong purifying selection in the core and a low number of pseudogenes

Universal and non-universal genes differ in the strength of purifying selection, with the nonsynonymous to synonymous substitutions ratio (dN/dS) being lower in the core and significantly higher in the periphery (Figure 5). This finding further supports the distinction between the highly conserved core and the evolvable periphery.

**Figure 5:**
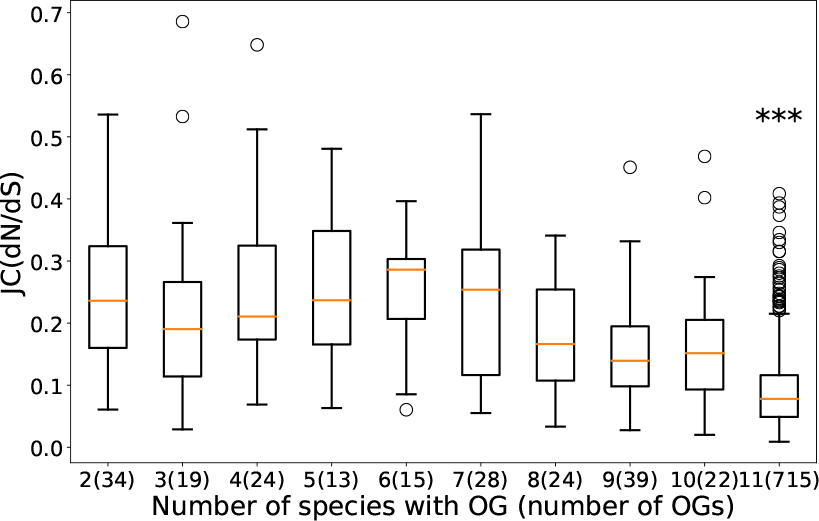
Nonsynonymous to synonymous substitutions ratio (dN/dS) as dependent on the number of species that have representatives of an Orthologous Group (OG) Only OGs with the median protein length above 100 amino acids are considered. Multiple substitutions per site are allowed (the Jukes–Cantor distance correction). The difference between medians is estimated with the nonparametric Mann–Whitney U test with the Bonferroni correction for multiple testing. *** denotes p > 0.001.

The number of genes with predicted frameshifts and internal stop codons in each individual genome is rather low, although it varies across species (on average, 11 in *C. trachomatis* vs. 35 in *C. psittaci*). In total, 140 OGs contain genes with frameshifts and nonsense mutations, including 82 OGs in the core. In particular, pseudogenes are present in such pathways as the TCA cycle, nucleotide metabolism, and homologous recombination. Several pathways exhibit parallel elimination of functionally related genes across the genus. Moreover, some genes in the population have been pseudogenised independently, as indicated by multiple frameshifts and internal stop codons in different parts of the phylogenetic tree. An interesting example is pseudogenisation in the TCA cycle across *C. trachomatis*, where multiple frameshifts and internal stop codons are identified in fumarate hydratase, class II (nonsense mutation in all trachoma strains (serovars A–C), frameshift in all LGV strains (L1–L3), succinate dehydrogenase flavoprotein (frameshift in some urogenital strains (D–K), nonsense mutation in some trachoma strains), succinate dehydrogenase iron-sulfur protein (nonsense mutation in all *C. trachomatis* strains), and succinate dehydrogenase cytochrome b558 subunit (nonsense mutation in most strains). All these genes are single-copy ones; frameshifts and nonsense mutations are at least sixty nucleotides upstream the gene end. Therefore, these cases are likely to represent true loss of function.

Most frameshifts and nonsense mutations seem to be rather recent, as none of them is present in the whole genus, and only few are completely species-specific. There are indications of a reduced purifying selection in OGs with high numbers of pseudogenes but we do not have sufficient power to estimate the effect due to the low number of such OGs (Supplementary file Figure 4).

### The chlamydial genomic diversity is mainly contained within several gene families and results from paralogisation

The majority of non-universal orthologous groups appear to be phylogenetically related and cluster into several broad orthologous families (OFs). The largest OFs are annotated as polymorphic membrane proteins, phospholipase D, ABC transporters, and multiple groups of inclusion membrane proteins (Table 2). Most OGs within families vary dramatically in terms of protein lengths and distribution across the genus, reflecting the processes of paralogisation, functional divergence, as well as pseudogenisation.

**Table 2:** Main families of orthologous groups (OFs). Only families with more than ten members are listed.

| OF id | Annotation | Number of OG in OF | Number of strains with OG (distribution) | Number of genes in OG (group size) | Median OG length, in aa |
| --- | --- | --- | --- | --- | --- |
| OF1000 | Polymorphic Outer Membrane proteins | 40 | 1 to 127 | 1 to 310 | 41 to 1752 |
| OF1001 | Inclusion membrane proteins | 20 | 1 to 21 | 1 to 21 | 40 to 549 |
| OF1002 | ABC transporter, ATP-binding proteins | 15 | 33 to 127 | 33 to 127 | 225 to 646 |
| OF1003 | Phospholipase D superfamily proteins | 15 | 1 to 127 | 1 to 127 | 6 to 420 |

Inclusion membrane proteins (Inc) are unique to phylum *Chlamydiae* and have been the object of extensive and comprehensive research (Heinz et al., 2010; Mital et al., 2010; Vorimore et al., 2013; Dehoux et al., 2011). Incs do not form a single gene family and are distinguished by their localization within the inclusion membrane rather than by sequence similarity. Here, Inc-like OGs have been defined as members of orthologous groups similar in sequence to annotated Incs. This resulted in 37 orthologous families comprised of two to twenty OGs each, and 144 individual OGs. The number of incs-like genes per genome varies from about 60 in *C. trachomatis* to 140 in *C. pneumoniae*, and they account for 6% and 12% of the total coding capacity in the respective species. The average length of an Inc-like protein is about 265 amino acids with the length range from 40 to 549 amino acids.

Phospholipase D (PLD) is another virulence-related, expanded gene family in *Chlamydia* (Coutinho-Silva et al., 2003; Nelson et al., 2006; Sait et al., 2014). All *Chlamydia* have two universally conserved genes containing the PLD-like domain (see Methods). One of these genes (assumed to encode cardiolipin synthase) is present as a single copy per genome, while the other one has multiple paralogs in the plasticity zone of *C. trachomatis*, *C. muridarum*, and *C. pecorum*, but not in other species. The plasticity zone complement of phospholipase D genes has been shown to represent an important strain-specific virulence factor (Nelson et al., 2006). Most chlamydial PLD-like genes encode 300-400 amino-acid proteins; however, short reading frames of varying length of 60-200 amino acids have also been identified.

Another important group of specific chlamydial proteins are Polymorphic membrane proteins (Pmp), which have been implicated in the host-cell attachment (Nunes et al., 2015; Kari et al., 2014; Tan et al., 2010; Wheelhouse et al., 2012; Gomes et al., 2006). The genes are all homologs and have been shown to be vertically inherited and conserved among *Chlamydiae*, *Verrucomicrobia*, *Lentisphaerae*, and *Planctomycetes* (Heinz et al., 2009). Here, all Pmp-like OGs (referred to as Pmps for simplicity, see Methods) formed a single OF comprised of 40 OGs. The average number of Pmps per genome varies significantly across the genus (on average, nine in *C. trachomatis* and *C. muridarum* vs. 23 in *C. pneumoniae* and 27 in *C. ibidis*). In total, they account for about 3% to 6% of the overall coding genomic capacity dependent on a strain. The average Pmp length is about 600 amino acids, while the overall range of lengths across Pmp-like genes is from 41 to 1752 amino acids. Functional Pmps are characterized by the domain structure comprised of the autotransporter domain (PF03797), the middle domain (PF07548), and at least one copy of the repeat domain (PF02415). However, the Pmp OF also contains genes missing some of these domains including multiple short Pmp-like genes (Figure 6).

**Figure 6:**
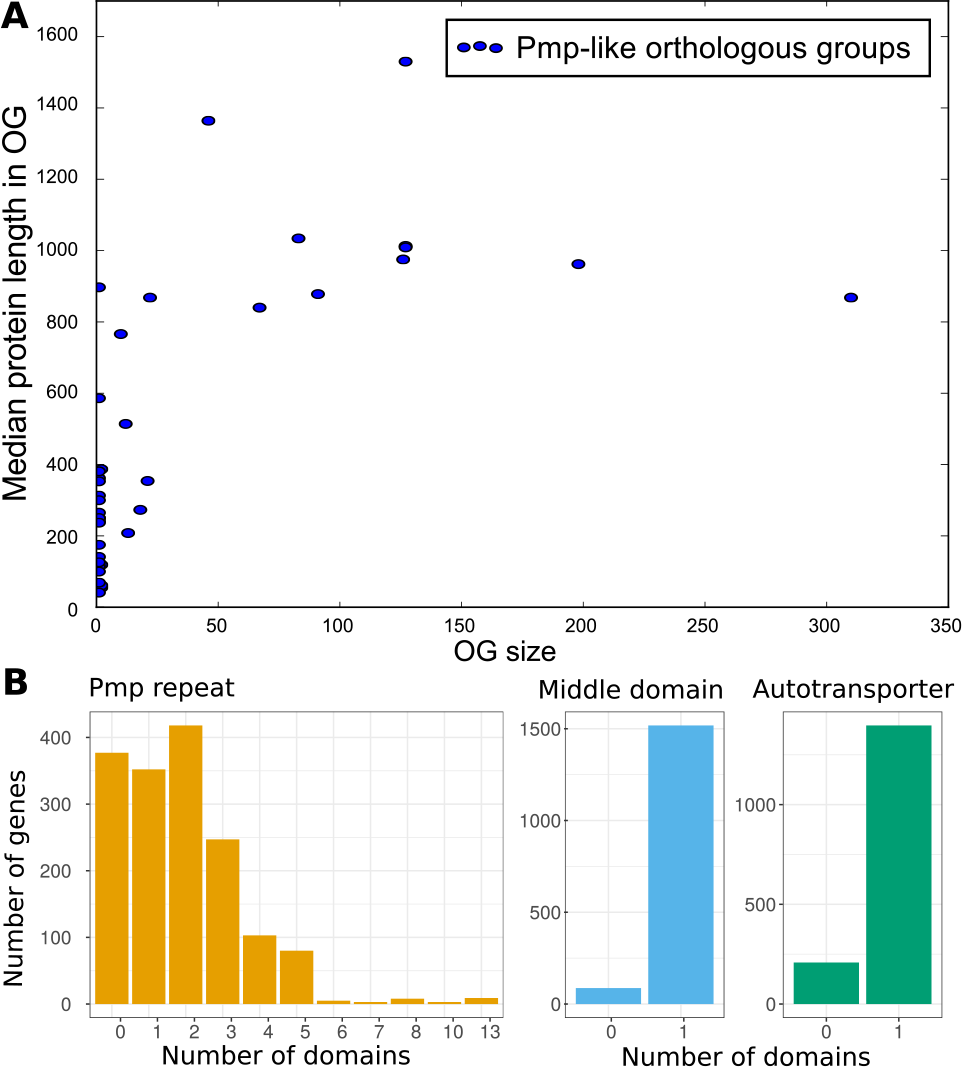
(A) Distribution of Orthologous Group (OG) sizes and median protein lengths within the OGs of Polymorphic outer membrane proteins (Pmps). (B) Distribution of the number of domains across Pmps.

Finally, many singletons and rare OGs are in fact short ORFs paralogous to core or species-specific genes. For example, one OF contains a universally conserved gene encoding succinate dehydrogenase cytochrome b558 subunit (290 amino acids long, OG1694), its shortened version presents in two strains (190 amino acids, OG2840), and one singleton (39 amino acids, S624).

### The PmpG subfamily of polymorphic membrane proteins demonstrates a combination of phase variation, gene duplication, and pseudogenisation processes

The largest OG in our analysis (OG1000) contained a subset of the pmpG subfamily of polymorphic membrane proteins (Grimwoodg and Stephens, 1999) in seven chlamydial species: *C. pneumoniae*, *C. psittaci*, *C. abortus*, *C. caviae*, *C. pecorum*, *C. ibidis*, and *C. gallinacea*. For simplicity, we annotated this OG as PmpG, keeping in mind that it also contains other Pmps with high sequence similarity to annotated PmpGs.

The entire family of Pmp proteins is related to virulence, and the PmpG subgroup has been shown to act as adhesins *in vitro* (Wehrl et al., 2004). The subset of pmpG genes clustered into OG1000 is of particular interest because it contains described cases of frameshifts due to the length variation of polyG tracts in *C. pneumoniae* (Pedersen, Christiansen, and Birkelund, 2001) and *C. abortus* (Wheelhouse et al., 2012). Replication-coupled frameshifting in polyG tracts (slipped strand mispairing) is considered to be a mechanism of phase variation, high-frequency on/off switching of phenotype expression involved in host adaptation and immune evasion (Henderson, Owen, and Nataro, 1999). In addition to the cases described above, PmpG orthologs with a broad range of polyG lengths (here we consider only polyG tracts of length 8 nucleotides or more) and multiple frameshifts at the homopolymeric tracts are present in *C. psittaci*, *C. caviae*, and *C. felis* suggesting that the phase variation mechanism is also active in these species. Furthermore, some pmpG genes in *C. pneumoniae* contain a polyC tract instead of the polyG one, and the former might also be subject to phase variation. The phylogenetic tree (Figure 7) shows that genes with polyG tracts in *C. pneumoniae* have diverged from the rest of polyG-containing genes in the other species, which could be explained either by two independent gains of the polyG tract (and presumably the phase variation capacity) or at least six independent losses.

**Figure 7:**
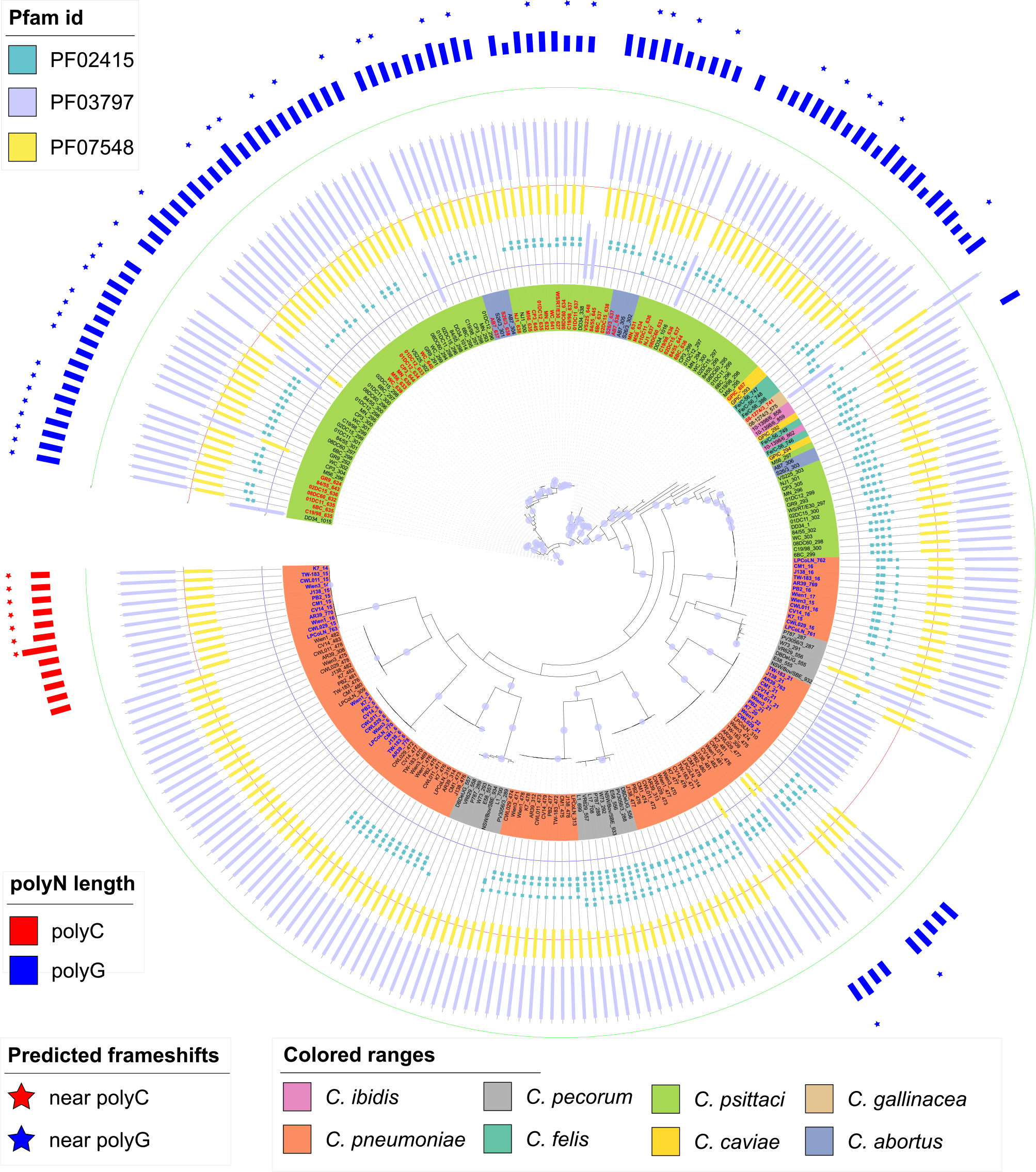
Phylogenetic tree for OG1000 (a subset of the pmpG group of Polymorphic membrane proteins). The label color on leaves corresponds to a genomic locus. PolyG/C tracts are defined as uninterrupted stretches of 8 or more guanines or cytosines, respectively. Frameshifts are shown on the leaves if they were predicted to occur at a distance not exceeding 30 nucleotides from the polyG/C tracts of the respective genes.

Our analysis also shows extensive paralogisation and pseudogenisation within OG1000, reflecting reduced evolutionary constraints on this group of genes. The number of paralogs per genome ranges from three to twelve with varying diversity within species (3-8 genes per strain in *C. psittaci* versus 10-12 genes per strain in *C. pneumonia*). These genes form one common chromosome cluster present in all seven species, and three smaller species-specific loci. In total, 50 out of 310 encoded proteins lack the autotransporter domain. Other genes are shorter than expected due to missing Pmp repeats or both repeats and the Pmp middle domain. Finally, the OG representatives feature an extremely high number of frameshifts and internal stop codons: 118 events in total with none to nine events per individual gene.

PolyG-containing pmpG genes are especially expanded in *C. psittaci*. The time of expansion is probably quite recent given very short branches in the phylogeneic tree. These genes do not cluster either by the presence of frameshifts or the length of polyG tracts, which agrees with the phase variation model. In addition, the analysis of full-length alignments of several subsets of *C. psittaci* OG1000 genes revealed multiple cases of short insertions and deletions resulting in frameshifts outside the polyG tract (data not shown). As a result, in some cases identical lengths of polyG tract correspond to a normal gene in some strains and a frameshifted one in others. These cases cannot be explained by the phase variation alone, and hence suggest additional mechanisms like gene conversion combined with degradation of pseudogenised paralogs.

PmpGs of *C. trahomatis* cluster in a distinct OG with one copy per genome. These genes also contain polyG tract of varying length, though all are in frame. In *in vitro* setups, the percentage of *C. trachomatis* inclusions not expressing pmpG varied from 1% to 10% in several independent experiments performed under the same infection conditions (Tan et al., 2010), hence suggesting a phase variation mechanism with frameshifting happening in a minor fraction of the clone populations. On the contrary, OG1000 was found to be the only Pmp-like OG in *Chlamydia* featuring multiple independent frameshifts along with high variation of polyG/C lengths and extensive paralogisation suggestive of a combination of phase variation and gene conversion mechanisms.

Finally, we checked how frequently homopolymeric tract occur in gene bodies of *Chlamydia*. Overall, homopolymeric tracts of length eight or more were present in about 15% of the chlamydial pan-genome, with 983 OGs containing at least one protein with a long homopolymeric tract. Long polyG and polyC tracts are not underrepresented in coding sequences of *Chlamydia* based on the Bernoulli model, unlike the situation in many other bacteria (Orsi, Bowen, and Wiedmann, 2009). OGs with genes carrying polyG/C tracts include chlamydial cytotoxin proteins, multiple groups of Pmps, Incs, phospholipase D, and hypothetical proteins suggesting possible phase variation. However, according to our data, frameshifts in polyN tracts are rare in the sequenced *Chlamydia* genomes and, in a number of cases, frameshifts in polyG/C tracts are single events probably representing true pseudogenes or sequencing errors.

## Discussion

The pan-genome of obligate intracellular bacteria of genus *Chlamydia* is characterized by a large pool of universally conserved genes, a small periphery, and relatively few strain-specific genes. The core, mainly comprised of the genes responsible for the information-processing machinery and central metabolism, evolves under strong purifying selection resulting in an extremely high conservation of sequences and of genomic locations. The periphery, on the other hand, is mostly species-specific, virulence-related, and evolving under reduced purifying selection. A subset of peripheral genes shows extensive length variations, domain losses, and multiple cases of independent frameshifting and gene losses in the population.

Several sources of evidence (saturation curves, paralogous families, analysis of gene losses and rearrangements) suggest that the majority of novel genes appear via paralogisation and functional divergence within just a few expanded gene families. Such confined pool for genomic innovations agrees with the isolated, intracellular lifestyle of *Chlamydia* and distinguishes them from the majority of prokaryotes, in which horizontal gene transfer is known to be one of the major drivers of genome evolution (Kunin and Ouzounis, 2003; Ochman, 2002). As an example, *Streptococcus pneumoniae* Shelyakin et al., 2018 or *Escherichia coli* (Gordienko, Kazanov, and Gelfand, 2013) have open pan-genomes where each new strain adds dozens of new genes, and up to 10% of periphery genes are spreading horizontally. Generally, pan-genomes with a large periphery are characteristic of organisms with large long-term effective population sizes and an ability to fill a variety of new niches (McInerney, McNally, and O’Connell, 2017).

Previous studies revealed widespread homologous recombination within chlamydial species (Gomes et al., 2007; DeMars and Weinfurter, 2008; Jeffrey et al., 2013; Joseph et al., 2012; Read et al., 2013). In this regard, an interesting research direction would be to assess whether homologous recombination contributes to further genomic innovations within expanded gene families and whether it counteracts accumulation of deleterious mutations in the core genes. Our analysis has identified several cases, where frameshifts at the same location in a gene are polyphyletically distributed in the gene’s phylogenetic tree, suggesting non-vertical inheritance. Similarly, multiple cases of short insertions and deletions have been found to result in frameshifts outside the polyG tract in the OG1000 genes. These cases might result from homologous recombination between paralogous gene copies within the same genome or between diverged orthologs in different strains. This analysis, however, is statistically underpowered by high sequence similarity within OGs and low bootstrap values of individual gene trees.

## Methods

### Data selection and pre-processing

A total of 146 genomes of *Chlamydia*, the only genus in family *Chlamydiaceae* according to a recently proposed classification (Stephens et al., 2009), were downloaded from NCBI Genbank (Benson et al., 2018). The sample included all complete genomes, as well as draft genomes assembled into at most 10 contigs, available as of June 5th, 2016. We excluded genomes sequenced after an in vitro recombination experiment (Jeffrey et al., 2013) and genomes with reported assembly anomalies (according to the NCBI Assembly webpage). If two genomes were assigned to the same strain or were identical based on the concatenated sequence of all universally conserved genes within the genus, only one genome from a pair was retained. The final sample was comprised of 115 complete and 12 draft genomes of genus *Chlamydia*. *Waddlia chondrophila* WSU 861044 (NC_014225) was selected as the outgroup to root the phylogenetic tree of the genus (Supplementary file Table 2). The final sample covered all known species of *Chlamydia* with the exception of *C. suis* due to the absence of a high quality genome at the time of analysis (Donati et al., 2014).

The analysis of chlamydial plasmids was beyond the scope of this study. However, some draft genome assemblies contained plasmid sequences as separate contigs. To exclude them, we performed BLASTn (Altschul et al., 1990) (the megablast algorithm) on all genomes using the set of known chlamydial plasmids as the database (accession numbers KT352924, NZ_CP009926, NC_002182, NC_018640, NZ_CM002267, AZNB01000165, CP006572, CP015841 and NC_007900) with the e-value threshold 10^3^. All contigs that had sequence identity of at least 98% and coverage along the contig length of at least 98% with at least one plasmid from the dataset were discarded.

The robustness of the chlamydial pan-genome definition relative to the sample size was estimated by reducing the number of genomes for species with many genomes. This influenced about 5% of OGs, mostly from paralogous gene families.

### Genome annotation

To exclude biases arising from differences in gene prediction algorithms, genes in all selected genomes were re-annotated de novo using the RAST pipeline with the “fix frameshifts” gene caller option (Overbeek et al., 2014). Gene functions were assigned by RAST. For all predicted CDSes in the genomes, additional domain search was performed with InterProScan-5.18-58.0 (Jones et al., 2014) (local installation, default parameters) using fifteen databases: SUPER-FAMILY v.1.75, Gene3D v.3.5.0, Hamap v.201605.11, Coils v.2.2.1, ProSiteProfiles v.20.119, ProSitePatterns v.20.119, TIGRFAM v.15.0, SMART v.7.1, CDD v.3.14, PRINTS v.42.0, PIRSF v.3.01, Pfam v.29.0, ProDom v.2006.1, PANTHER v.10.0, and InterPro patterns database v.58.0. Assignment of the Enzyme Commission (EC) numbers and the Gene Onthology (GO) terms to individual genes was included in the RAST annotation pipeline. In addition, GO-terms were assigned to individual Pfam domains by InterProScan.

### Building the pan-genome: Orthologous Groups and Orthologous Families of genes

Orthologous groups (OGs) of genes were identified using the orthoMCL v.2.0.9 software package (default parameters, e-value cutoff of 10^5^, mcl algorithm inflation value 1.5)(Chen et al., 2006). Clustering the full complement of protein-coding chlamydial genes yielded a set of 2595 orthologous groups (OGs) of genes further referred to as the pan-genome. This set was subdivided into universally conserved genes present in every genome in the sample (core), genes present in some but not all strains (periphery), and unique strain-specific genes (singletons). A standard accession identifier was assigned to each OG based on the best BLAST hit against the RefSeq database (O’Leary et al., 2016) (coverage above 50 amino acids, other parameters set to default). In addition, the encoded proteins comprising core, periphery, and singletons were functionally annotated using RAST and InterProScan software (Aziz et al., 2008; Jones et al., 2014). A function was assigned to each OG based on the most frequent predicted function of its members in the RAST annotation.

More distant relationships among OGs were determined by an additional round of OrthoMCL applied to one representative from each OG (with the gene length closest to the group median). As a result, 508 orthologous groups were grouped into 157 clusters called orthologous families (OFs). In total, 27 OFs contained four or more OGs (Supplementary file Table 3, the og2of column).

### Pan-genome annotation

OGs were assigned to Clusters of Orthologous Genes (COG) using the COGnitor software (Galperin et al., 2015). Assignment to KEGG (Kanehisa et al., 2016) pathways was performed using the BlastKOALA software (Kanehisa, Sato, and Morishima, 2016). GO enrichment analysis was done using R GSEABase and GOstats packages (Falcon and Gentleman, 2007).

Inclusion membrane proteins (Incs) were annotated based on sequence similarity to previously identified Incs (Dehoux et al., 2011). Amino acid sequences for the list of 461 Incs annotated in six chlamydial species (*C. trachomatis*, *C. pneumoniae*, *C. felis*, *C. abortus*, *C. caviae*, and *C. muridarum*) were downloaded from NCBI Genbank and scanned against representatives of OGs using BLASTp (e-value cutoff 0.01, coverage cutoff 50 amino acids). As the result, 285 Orthologous Groups were annotated as “Inc-like”. However, some Incs may have been missed, because genes unique to the remaining five species could not be considered. De novo annotation of Incs requires recognition of a specific secondary structure motif, which is beyond the scope of this study.

Polymorphic Outer Membrane proteins (Pmps) were initially annotated based on the presence of characteristic Pfam domains PF03797 (Autotransporter domain), PF07548 (*Chlamydia* polymorphic membrane protein middle domain), and PF02415 (*Chlamydia* polymorphic membrane protein repeat) (Wehrl et al., 2004; Wheelhouse et al., 2012; Pedersen, Christiansen, and Birkelund, 2001; Oomen et al., 2004). All Pmp genes were found to be homologous and to fall within one orthologous family (OF1000). It should be noted that this OF also contains many shorter genes, lacking some or all characteristic Pmp domains, which probably are pseudogenes. All genes falling into OF1000 were annotated as “Pmp-like”.

Phospholipase D endonuclease superfamily (PLD) proteins were annotated based on the presence of PLD-like domain PF13091 (Ponting and Kerr, 1996). This domain was identified in all genes from one distinct orthologous group (OG1391) and in the majority of genes belonging to a broad family of 15 OGs (OF1003). All these OGs were annotated as “PLD-like”.

The resulting annotation of orthologous groups is provided in Supplementary file Table 2.

### Draft genomes and short reading frames

The inclusion of draft genomes did not distort the shape of the distribution of orthologous groups by the number of strains (Supplementary file Figure 5). The number of orthologous groups universally present in the whole genus reduced from 729 to 725 after including twelve draft genomes into the analysis. These four OGs were absent exclusively in *Chlamydia ibidis*, which was represented in Genbank by a single draft genome. Therefore, no sequencing artefacts were likely to be introduced in our pan-genome analysis by including draft genomes.

The analysis of genomic rearrangements and gene upstream regions was done for the complete genomes only to avoid distortions due to incomplete assembly.

Some artifacts in the prediction of short reading frames might have been introduced by the annotation pipeline (Ochman, 2002). Out of 1370 OGs with median protein length below 100 amino acids, 352 groups had no homologs in the RefSeq database; three more OGs without RefSeq hits had lengths between 100 and 200 amino acids. On the other hand, at least one small protein (9kDa OmcA) was not predicted to be encoded in the *Chlamydia pneumoniae* genomes by the RAST annotation pipeline, though the corresponding open reading frames could be identified by tBLASTn. Most of these short OGs had no functional annotation (1302 OGs) and could not be assigned to any known COG (1304 OGs). Furthermore, out of these 1370 OG only 30 were universally conserved across the whole genus (mainly, ribosomal proteins). However, short genes (protein length below 100 amino acids) were included only into the general pan-genome statistics, as no in-depth analysis could be done for them without additional data.

In general, our predicted pan-genome and coregenome sizes agree well with the previously described data (Psomopoulos et al., 2012; Collingro et al., 2011), and therefore no large biases are expected to be introduced into the analysis during genome annotations.

### Prediction of pseudogenes

Frameshifts and internal stop codons were predicted within the RAST annotation pipeline by enabling the “fix frameshifts” option, and the coordinates of frameshifts were defined relative to the gene end. The distribution of these events was found to be nearly uniform along the gene length (*p*=0.5906, KolmogorovSmirnov test), therefore we could not set an obvious cut-off for non-deleterious events. For the purpose of functional analysis, positions of frameshifts or internal stop codons relative to the gene end as well as their occurrence relative to predicted protein domains were considered in each case individually. For the pN/pS analysis (Suppl Figure S3), all OGs with the median protein length above 100 amino acids and predicted frameshifts or internal stop codons not less than 60 nucleotides from the end were considered.

In the original algorithm, this option is mainly targeted at detecting frameshifts caused by sequencing errors (Aziz et al., 2008). However, since it is highly unlikely to get an error at exactly the same nucleotide position in two independent sequencing runs, we treated such cases as true pseudogenisation events. With this approach, we are limited to detecting frameshifts in species represented by at least two genomes, but this helps to avoid potential false positives. The only exception was made for the analysis of frameshifts near polyN tracts. Since these sequences are known to be mutagenic (Belkum et al., 1998), and pmp genes truncated at polyG tracts have been detected in *Chlamydia* in in vivo experiments (Pedersen, Christiansen, and Birkelund, 2001; Wheelhouse et al., 2012), there events were considered separately.

### Homopolymeric tracts

The expected frequencies of homopolymeric tracts in coding sequences were calculated using the Bernoulli model with observed frequencies of nucleotides using formulae given in (Bizzaro and Marx, 2003). In the OG1000 analysis, we focused only on homopolymeric tracts of length eight or more nucleotides (based on cases described in the literature) and did not allow substitutions within the stretches. Therefore, the absence of a polyG tract might also mean shortened or mutated homopolymeric sequences.

### Alignments and phylogenetic analysis

Nucleotide and protein alignments were constructed using the MACSE (Ranwez et al., 2011) and MUSCLE algorithms (Edgar, 2004), respectively. The phylogeny was reconstructed using the RaxML toolkit (GTRGAMMA and PROTGAMMABLOSUM62 models, 100 bootstrap replicates). For an unrooted phylogenetic tree, we used the concatenated nucleotide sequence of 446 genes that were shared by all 127 strains, were present as one copy per genome, and contained no predicted frameshifts or nonsense mutations. For a rooted tree, we added a requirement for universal genes to be also shared (single copy per genome, no predicted frameshifts or nonsense mutations) with the outgroup, *Waddlia chondrophila* WSU 86-1044; thus only 346 genes remained. Phylogenetic trees were visualized using the iTOL online tool (Letunic and Bork, 2016). The resulting phylogenetic tree is provided as Supplementary file Figure 6.

### Reconstruction of genomic rearrangements

Stable common gene blocks, i.e. groups of common genes that maintain the same order across all considered genomes, were reconstructed using the DRIMM-Synteny algorithm (Pham and Pevzner, 2010) for 115 complete genomes and 715 single-copy genes universally conserved in these genomes. Evolutionary history of inversions was reconstructed by MGRA 2.2 (Avdeyev et al., 2016) using the rooted phylogenetic tree built as described above.

To detect the inflow of genes transferred horizontally, we used the following model. If a gene was inherited from the common ancestor and it was lost by several of the succeeding strains, we expected to detect it at the same locus in the remaining strains. On the contrary, genes transferred horizontally may occupy in different loci in different strains. Positions of non-universal genes, to compensate for their variability, were defined relative to the nearest universal genes. Genes whose neighboring universal genes were affected by rearrangements (according to our reconstruction), i.e. genes located at or near boundaries of stable common gene blocks, were filtered out.

### dN/dS and pN/pS calculations

To estimate nonsynonymous to synonymous substitution ratio between (dN/dS) or within (pN/pS) species, we used the KaKs_Calculator Toolbox version 2.0 with default parameters (Zhang et al., 2006). Multiple substitutions were allowed by applying the Jukes-Cantor correction (Jukes and Cantor, 1969). Only orthologous groups with the median length exceeding 100 amino acids and containing no paralogs were considered.

The analysis of the pN and pS values covered species represented by at least two strains. Calculations were done for each pair of strains within species, and the median pN/pS ratio was assigned to respective OGs. For the dN and dS calculations, we performed 30 rounds of random selection of pairs of strains from two different species. The median dN/dS ratio was then assigned to respective OGs.

## Declarations

### Availability of data and materials

All datasets on which the conclusions of the paper rely presented in the main manuscript and additional supporting files.

### Author’s contributions

OS, AVC and MSG conceived the general approach of the study, OS, AVC, OOB, PS, VAF, EA and VB performed a data analysis; OS, AVC, OOB and MSG wrote the paper.

### Ethics approval and consent to participate

Not applicable

### Consent for publication

Not applicable

### Additional Files

**Supplementary file Fig. S1.** The number of new genes added to the pangenome upon addition of new strains *Chlamydia* spp. (a) for all genes (b) with exclusion of short reading frames. The number of new genes is plotted as a function of the number (*n*) of strains sequentially added (see the model in Donati et al., 2010). For each *n*, points are the values obtained for different strain combinations; red symbols are the averages of these values. The superimposed line is the best fit with a decaying power law y *y* = *A*^B^.

**Supplementary file Fig. S2.** Genes with mosaic phyletic pattern on the phylogenetic tree of the genus. Each column is an orthologous group, star indicates presence of the OG in a genome. Genes that changed their genomic context in some strains are shown in red.

**Supplementary file Fig. S3.** Phylogenetic tree of the orthologous group OG 2167 coded an outer membrane protein revealing multiple, independent gene transfers into different strains.

**Supplementary file Fig. S4.** Nonsynonymous to synonymous substitutions ratio (dN/dS) depending on the presence and number of genes with frameshift or internal stop codons in the OG. Numbers of corresponding OGs are shown in brackets. Only OGs with the median protein length above 100 amino acids are considered. Multiple substitutions per site are allowed (the Jukes–Cantor distance correction). The difference between medians is estimated with the nonparametric Mann–Whitney U test with the Bonferroni correction for multiple testing. * denotes *p* < 0.05

**Supplementary file Fig. S5.** Distribution of Orthologous Groups (OGs) by their presence in different subsets of *Chlamydia* strains (the U-curve) based only on complete genomes (red) or including draft genomes (black).

**Supplementary file Fig. S6.** Phylogenetic tree of the genus *Chlamydia* based on concatenated nucleotide sequence of 446 genes shared by all 127 strains. Bootstrap values are shown on the edges. *Waddlia chondrophila* WSU 86-1044 is taken as an outgroup to root the tree.

**Supplementary file Table S1.** Summary of pangenome by species.

Ref: RefSeq genome ID; genome_id: internal genome ID; organism: full strain name; contigs: 1 for complete genome, otherwise number of contains; gene_number: number of genes predicted by RAST pipeline (see Methods); DNA length: genome length in nucleotides.

**Supplementary file Table S2.** Summary of the analysed genomes.

Group id: internal OG ID; Product: most frequent annotation of proteins from the OG; Og2Of: if OG was signed to prthologous family, corresponding OF ID (see Methods); Representative sequence: amino acid sequence of a single representative of OG with the length closest to the group median.

**Supplementary file Table S3.** Summary of orthologous groups.

## Bibliography

AbdelRahman, YM and RJ Belland (2005). “The chlamydial developmental cycle”. In: FEMS Microbiology Reviews 29.5, pp. 949–959.

Altschul, SF et al. (1990). “Basic local alignment search tool”. In: Journal of Molecular Biology 215.3, pp. 403–410

Avdeyev, P et al. (2016). “Reconstruction of ancestral genomes in presence of gene gain and loss”. In: Journal of Computational Biology 23.3, pp. 150–164.

Aziz, RK et al. (2008). “The RAST Server: rapid annotations using subsystems technology”. In: BMC Genomics 9, p. 75.

Bachmann, NL, A Polkinghorne, and P Timms (2014). “Chlamydia genomics: providing novel insights into chlamydial biology”. In: Trends in Microbiology 22.8, pp. 464–472.

Barry, CE 3rd, SF Hayes, and T Hackstadt (1992). “Nucleoid condensation in Escherichia Coli that express a chlamydial histone homolog”. In: Science 256.5055, 377–79.

Belkum, A van et al. (1998). “Short-sequence DNA repeats in prokaryotic genomes”. In: Microbiol Mol Biol Rev 62.2, pp. 275–93.

Belland, RJ et al. (2001). “Chlamydia trachomatis cytotoxicity associated with complete and partial cytotoxin genes”. In: Proc Natl Acad Sci U S A 98.24, pp. 13984–9.

Benson, DA et al. (2018). “GenBank”. In: Nucleic Acids Research 45.D1, pp. D41–D47.

Bizzaro, JW and KA Marx (2003). “Poly: a quantitative analysis tool for simple sequence repeat (SSR) tracts in DNA”. In: BMC Bioinformatics 4, p. 22.

Caldwell, HD et al. (2003). “Polymorphisms in Chlamydia trachomatis tryptophan synthase genes differentiate between genital and ocular isolates”. In: J Clin Invest. 111.11, pp. 1757–69.

Campbell, LA and ME Rosenfeld (2014). “Persistent C. pneumoniae infection in atherosclerotic lesions: rethinking the clinical trials.” In: Frontiers in Cellular and Infection Microbiology 4, p. 34.

Chao, A (1987). “Estimating the population size for capture-recapture data with unequal catchability.” In: Biometrics 43, pp. 783–91.

Chen, F et al. (2006). “OrthoMCL-DB: querying a comprehensive multi-species collection of ortholog groups.” In: Nucleic Acids Research 34 (Database issue), pp. D363–8.

Chumduri, C et al. (2013). “Chlamydia infection promotes host DNA damage and proliferation but impairs the DNA damage response.” In: Cell Host Microbe 13.6, pp. 746–58.

Collingro, A et al. (2011). “Unity in variety the pangenome of the Chlamydiae.” In: Mol Biol Evol. 28.12, pp. 3253–70.

Coutinho-Silva, R et al. (2003). “Inhibition of chlamydial infectious activity due to P2X7R-dependent phospholipase D activation.” In: Immunity 19.3, pp. 403–412.

Dehoux, P et al. (2011). “Multi-genome identification and characterization of chlamydiae-specific type III secretion substrates: the Inc proteins.” In: BMC Genomics 12, p. 109.

DeMars, R and J Weinfurter (2008). “Interstrain gene transfer in Chlamydia trachomatis in Vitro: mechanism and significance.” In: Journal of Bacteriology 190.5, 1605–14.

Domman, D, and M Horn (2015). “Following the foot-steps of chlamydial gene regulation.” In: Molecular Biology and Evolution 32.12, 3035–46.

Donati, C et al. (2010). “Structure and dynamics of the pan-genome of Streptococcus pneumoniae and closely related species.” In: Genome Biol. 11.10, R107.

Donati, M et al. (2014). “Genome sequence of Chlamydia Suis MD56, isolated from the conjunctiva of a weaned piglet.” In: Genome Announc 2.3, e00425–14

Doniger, J et al. (1992). “The product of Unr, the highly conserved gene upstream of N-Ras, contains multiple repeats similar to the cold-shock domain (CSD), a putative DNA-binding motif.” In: The New Biologist 4.4, 389–95.

ECDC (2015). Guidance on chlamydia control in Europe. doi:10.2900/667703. Stockholm.

Edgar, RC (2004). “MUSCLE: multiple sequence alignment with high accuracy and high throughput.” In: Nucleic acids research 32.5, pp. 1792–1797.

Elwell, C, K Mirrashidi, and J Engel (2016). “Chlamydia cell biology and pathogenesis.” In: Nature Reviews Microbiology 14.6, pp. 385–400.

Everett, KD, RM Bush, and AA. Andersen (1999). “Emended description of the order Chlamydiales, proposal of Parachlamydiaceae fam. nov. and Simkaniaceae fam. nov., each containing one monotypic genus, revised taxonomy of the family Chlamydiaceae, including a new genus and five new species, and standards for the identification of organisms.” In: Int J Syst Bacteriol. 49 Pt 2, pp. 415–40.

Falcon, S and R Gentleman (2007). “Using GOstats to test gene lists for GO term association.” In: Bioinformatics 23.2, 257–58.

Galperin, MY et al. (2015). “Expanded microbial genome coverage and improved protein family annotation in the COG database.” In: 43.Database issue, pp. D261–9.

Goldstein, J, NS Pollitt, and M Inouye (1990). “Major cold shock protein of Escherichia coli.” In: Proceedings of the National Academy of Sciences 87.1, pp. 283–287

Gomes, JP et al. (2006). “Polymorphisms in the nine polymorphic membrane proteins of Chlamydia trachomatis across all serovars: evidence for serovar Da recombination and correlation with tissue tropism.” In: Journal of Bacteriology 188.1, pp. 275–286.

Gomes, JP et al. (2007). “Evolution of Chlamydia trachomatis diversity occurs by widespread interstrain recombination involving hotspots.” In: Genome Research 17.1, pp. 50–60.

Gordienko, EN, MD Kazanov, and MS Gelfand (2013). “Evolution of pan-genomes of Escherichia coli, Shigella spp., and Salmonella enterica.” In: Journal of bacteriology 195.12, pp. 2786–2792.

Grimwoodg, J and RS Stephens (1999). “Computational analysis of the polymorphic membrane protein superfamily of Chlamydia trachomatis and Chlamydia pneumoniae.” In: Microbial & Comparative Genomics 4.3, pp. 187–201.

Heinz, E et al. (2009). “Comprehensive in silico prediction and analysis of chlamydial outer membrane proteins reflects evolution and life style of the Chlamy-diae.” In: vol. 10, p. 634.

Heinz, E et al. (2010). “Inclusion membrane proteins of Protochlamydia amoebophila UWE25 reveal a conserved mechanism for host cell interaction among the Chlamydiae.” In: Journal of bacteriology 192. 19, pp. 5093–102.

Henderson, I, P Owen, and J Nataro (1999). “Molecular switches — the ON and OFF of bacterial phase variation.” In: Molecular Microbiology 33, pp. 919–932

Jacquier, N, PH Viollier, and G Greub (2015). “The role of peptidoglycan in chlamydial cell division: towards resolving the chlamydial anomaly.” In: FEMS microbiology reviews 39. 2, pp. 262–75.

Jeffrey, BM et al. (2013). “Genomic and phenotypic characterization of in vitro-generated Chlamydia tra-chomatis recombinants.” In: vol. 13, p. 142.

Jones, P et al. (2014). “InterProScan 5: genome-scale protein function classification.” In: Bioinformatics 30.9, pp. 1236–1240.

Joseph, SJ et al. (2012). “Population genomics of Chlamydia trachomatis: insights on drift, selection, recombination, and population structure.” In: Molecular biology and evolution 29. 12, pp. 3933–46.

Jukes, TH and CR Cantor (1969). “Evolution of protein molecules.” In: chap. 24, pp. 21–132.

Kanehisa, M, Y Sato, and K Morishima (2016). “BlastKOALA and GhostKOALA: KEGG tools for functional characterization of genome and metagenome sequences.” In: Journal of molecular biology 428.4, pp. 726–731.

Kanehisa, M et al. (2016). “KEGG as a reference resource for gene and protein annotation.” In: Nucleic acids research 44.D1, pp. D457–62.

Kari, L et al. (2014). “Chlamydia trachomatis polymorphic membrane protein D is a virulence factor involved in early host-cell interactions.” In: Infection and immunity 82.7, pp. 2756–62.

Karunakaran, KP et al. (2003). “Molecular analysis of the multiple GroEL proteins of Chlamydiae.” In: Journal of bacteriology 185.6, pp. 1958–66.

Koonin, EV and YI Wolf (2008). “Genomics of bacteria and archaea: the emerging dynamic view of the prokaryotic world.” In: vol. 36. 21, pp. 6688–719.

Kunin, V and CA. Ouzounis (2003). “The balance of driving forces during genome evolution in prokaryotes.” In: Genome Res. 13. 7, pp. 1589–94.

Letunic, I and P Bork (2016). “Interactive tree of life (iTOL) v3: an online tool for the display and annotation of phylogenetic and other trees.” In:

Liechti, G et al. (2016). “Pathogenic Chlamydia lack a classical sacculus but synthesize a narrow, midcell peptidoglycan ring, regulated by MreB, for cell division.” In: PLoS Pathog. 12.5, e1005590.

McInerney, JO, A McNally, and MJ O’Connell (2017). “Why prokaryotes have pangenomes.” In: Nat Microbiol 2, p. 17404.

McNally, D and MA Fares (2007). “In silico identification of functional divergence between the multiple groEL gene paralogs in Chlamydiae.” In: BMC Evolutionary Biology 7, p. 81.

Mital, J et al. (2010). “Specific chlamydial inclusion membrane proteins associate with active Src family kinases in microdomains that interact with the host microtubule network.” In: Cellular microbiology 12.9, pp. 1235–49.

Moldovan, MA and MS Gelfand (2018). “Pangenomic definition of prokaryotic species and the phylogenetic structure of Prochlorococcus spp.” In: Front Microbiol. 9, p. 428.

Moore, ER and SP Ouellette (2014). “Reconceptualizing the chlamydial inclusion as a pathogen-specified parasitic organelle: an expanded role for Inc proteins.” In: vol. 4, p. 157.

Moran, NA (2002). “Microbial minimalism: genome reduction in bacterial pathogens.” In: Cell 108.5, pp. 583–586.

Nelson, DE et al. (2006). “Inhibition of chlamydiae by primary alcohols correlates with the strain-specific complement of plasticity zone phospholipase D genes.” In: Infection and immunity 74.1, pp. 73–80.

Nunes, A and JP Gomes (2014). “Evolution, phylogeny, and molecular epidemiology of Chlamydia.” In: Infection, Genetics and Evolution 23, pp. 49–64.

Nunes, A et al. (2015). “Bioinformatic analysis of Chlamydia trachomatis polymorphic membrane proteins PmpE, PmpF, PmpG and PmpH as potential vaccine antigens.” In: vol. 10. 7, e0131695.

Ochman, H (2002). “Distinguishing the ORFs from the ELFs: short bacterial genes and the annotation of genomes.” In: Trends Genet. 18.7, pp. 335–7.

O’Leary, NA et al. (2016). “Reference sequence (Ref-Seq) database at NCBI: current status, taxonomic expansion, and functional annotation.” In: vol. 44. D1, pp. D733–45.

Omsland, A et al. (2012). “Developmental stage-specific metabolic and transcriptional activity of Chlamydia trachomatis in an axenic medium.” In: Proceedings of the National Academy of Sciences of the United States of America 109.48, pp. 19781–5.

Omsland, A et al. (2014). “Chlamydial metabolism revisited: interspecies metabolic variability and developmental stage-specific physiologic activities.” In: FEMS Microbiol Rev. 38.4, pp. 779–801.

Oomen, CJ et al. (2004). “Structure of the translocator domain of a bacterial autotransporter.” In: EMBO J. 23.6, pp. 1257–66.

Orsi, RH, BM Bowen, and M Wiedmann (2009). “Homopolymeric tracts represent a general regulatory mechanism in prokaryotes.” In: vol. 11, p. 102.

Overbeek, RA et al. (2014). “The SEED and the rapid annotation of microbial genomes using subsystems technology (RAST).” In: vol. 42. Database issue, pp. D206–14.

Pedersen, AS, G Christiansen, and S Birkelund (2001). “Differential expression of Pmp10 in cell culture infected with Chlamydia pneumoniae CWL029.” In: FEMS microbiology letters 203.2, pp. 153–9.

Pham, SK and PA Pevzner (2010). “DRIMM-Synteny: decomposing genomes into evolutionary conserved segments.” In: Bioinformatics 26, pp. 2509–16.

Ponting, CP and ID Kerr (1996). “A novel family of phospholipase D homologues that includes phospholipid synthases and putative endonucleases: identification of duplicated repeats and potential active site residues.” In: Protein Sci. 5.5, pp. 914–22.

Psomopoulos, FE et al. (2012). “The chlamydiales pangenome revisited: structural stability and functional coherence.” In: vol. 3. 2, pp. 291–319.

Ranwez, V et al. (2011). “MACSE: Multiple Alignment of Coding SEquences accounting for frameshifts and stop codons.” In:

Read, TD et al. (2000). “Genome sequences of Chlamydia trachomatis MoPn and Chlamydia pneumoniae AR39.” In: Nucleic acids research 28.6, pp. 1397–406.

Read, TD et al. (2003). “Genome sequence of Chlamydophila caviae (Chlamydia psittaci GPIC): examining the role of niche-specific genes in the evolution of the Chlamydiaceae.” In: Nucleic acids research 31.8, pp. 2134–47.

Read, TD et al. (2013). “Comparative analysis of Chlamydia psittaci genomes reveals the recent emergence of a pathogenic lineage with a broad host range.” In: vol. 4. 2, e00604–12.

Ross, W et al. (2013). “The magic spot: a ppGpp binding site on E. coli RNA polymerase responsible for regulation of transcription initiation.” In: Molecular cell 50.3, pp. 420–9.

Rouli, L et al. (2015). “The bacterial pangenome as a new tool for analysing pathogenic bacteria.” In:

Sachse, K et al. (2014). “Evidence for the existence of two new members of the family Chlamydiaceae and proposal of Chlamydia avium sp. nov. and Chlamydia gallinacea sp. nov.” In: Systematic and applied microbiology 37.2, pp. 79–88.

Sachse, K et al. (2015). “Emendation of the family Chlamydiaceae: proposal of a single genus, Chlamydia, to include all currently recognized species.” In: Systematic and applied microbiology 38.2, pp. 99–103

Sait, M et al. (2014). “Genome sequencing and comparative analysis of three Chlamydia pecorum strains associated with different pathogenic outcomes.” In: BMC Genomics 15, p. 23.

Schwöppe, C, HH Winkler, and HE Neuhaus (2002). “Properties of the glucose-6-phosphate transporter from Chlamydia pneumoniae (HPTcp) and the glucose-6-phosphate sensor from Escherichia coli (UhpC).” In: J Bacteriol. 184.8, pp. 2108–15.

Shaw, EI et al. (2000). “Three temporal classes of gene expression during the Chlamydia trachomatis developmental cycle.” In: Molecular microbiology 37.4, pp. 913–25.

Shelyakin, Pavel V et al. (2018). “Comparative analysis of Streptococcus genomes.” In: BioRxiv. doi:10.1101/447938.

Stephens, RS et al. (2009). “Divergence without difference: phylogenetics and taxonomy of Chlamydia resolved.” In: FEMS immunology and medical microbiology 55.2, pp. 115–9.

Syal, K and D Chatterji (2015). “Differential binding of ppGpp and pppGpp to E. coli RNA polymerase: photo-labeling and mass spectral studies.” In: Genes to cells: devoted to molecular cellular mechanisms 20.12, pp. 1006–16.

Tan, CY et al. (2010). “Variable expression of surface-exposed polymorphic membrane proteins in in vitro-grown Chlamydia trachomatis.” In: Cellular microbiology 12.2, pp. 174–87.

Tettelin, H et al. (2005). “Genome analysis of multiple pathogenic isolates of Streptococcus agalactiae: implications for the microbial “pan-genome”.” In: Proceedings of the National Academy of Sciences of the United States of America 102.39, pp. 13950–5.

Tjaden, J et al. (1999). “Two nucleotide transport proteins in Chlamydia trachomatis, one for net nucleoside triphosphate uptake and the other for transport of energy.” In: Journal of bacteriology 181.4, pp. 1196–202.

Villareal, C, JA Whittum-Hudson, and AP Hudson (2002). “Persistent Chlamydiae and chronic arthritis.” In: Arthritis Res. 4.1, pp. 5–9.

Vorimore, F et al. (2013). “Isolation of a new Chlamydia species from the feral sacred ibis (Threskiornis aethiopicus): Chlamydia ibidis.” In:

Wehrl, W et al. (2004). “From the inside out– processing of the Chlamydial autotransporter PmpD and its role in bacterial adhesion and activation of human host cells.” In: Molecular microbiology 51.2, pp. 319–34.

Wheelhouse, N et al. (2012). “Expression patterns of five polymorphic membrane proteins during the Chlamydia abortus developmental cycle.” In:

Xu, L et al. (2006). “Average gene length is highly conserved in prokaryotes and eukaryotes and diverges only between the two kingdoms.” In: Mol Biol Evol. 23.6, pp. 1107–1108.

Zhang, Z et al. (2006). “KaKs_Calculator: calculating Ka and Ks through model selection and model averaging.” In: Genomics Proteomics Bioinformatics 4.4, pp. 259–263.

