## Supplementary figures and images for "*Chlamydia* pan-genomic analysis reveals balance between host adaptation and selective pressure to genome reduction"

### Supplemental file Figure 1

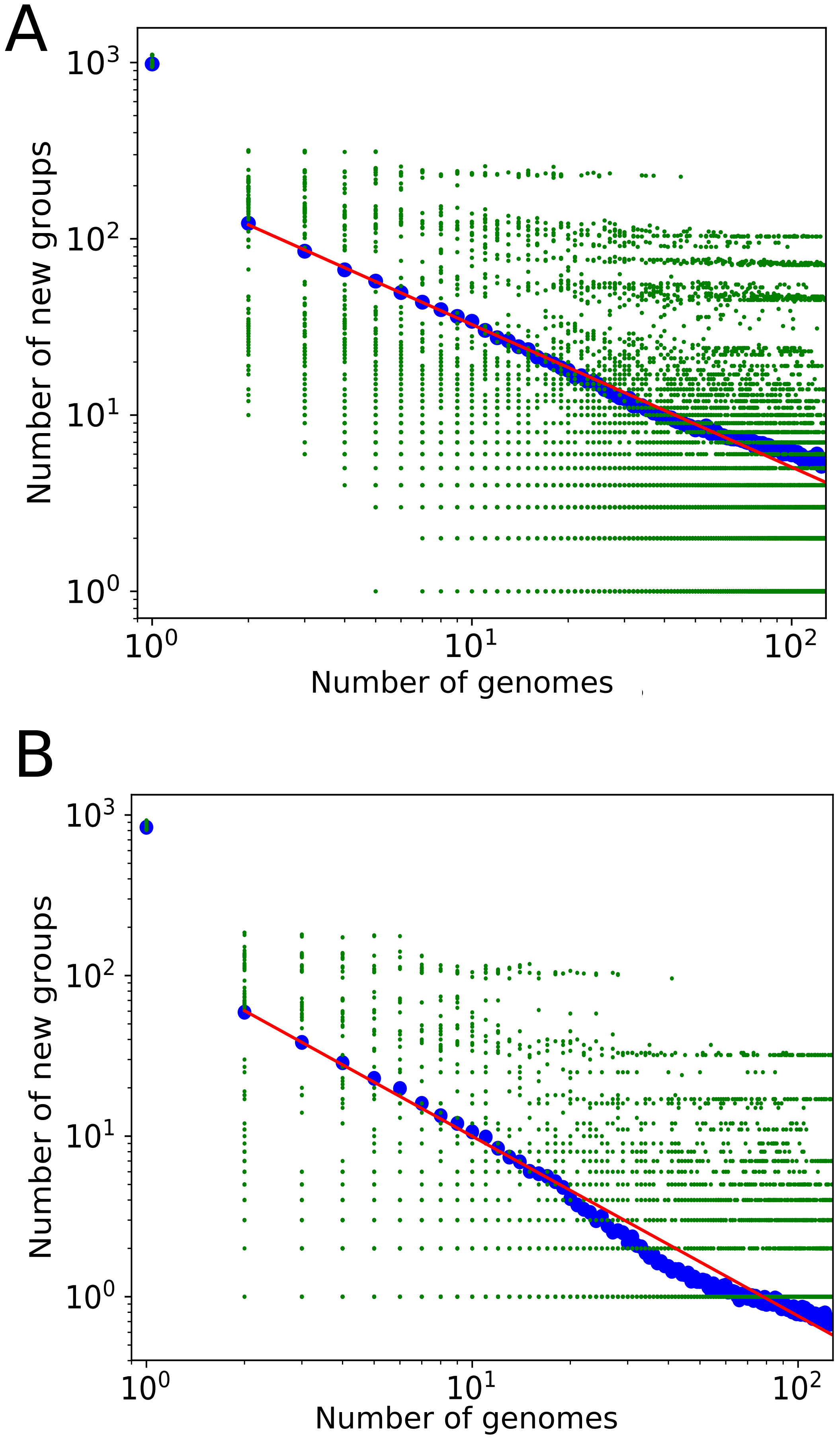

### Supplemental file Figure 2

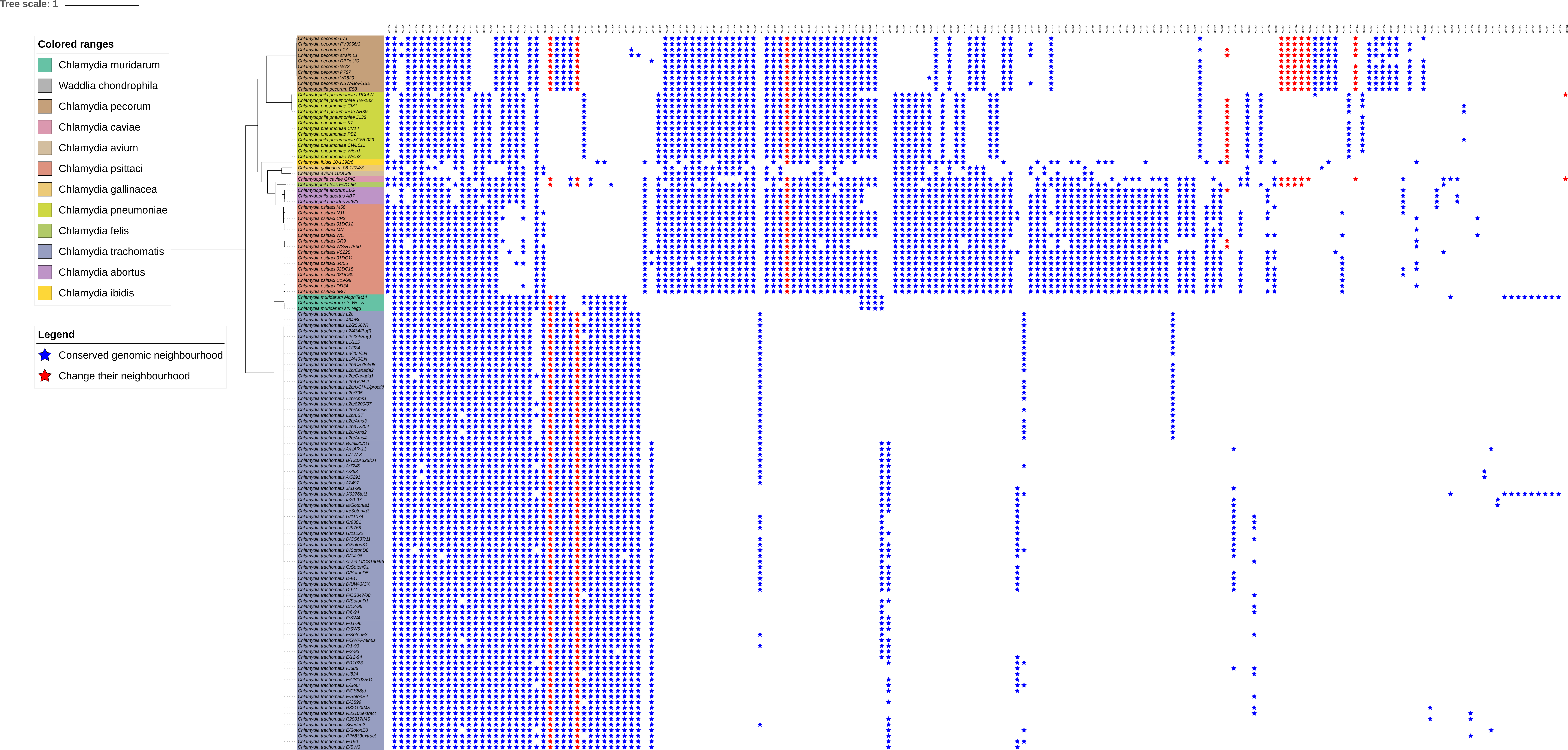

### Supplemental file Figure 3

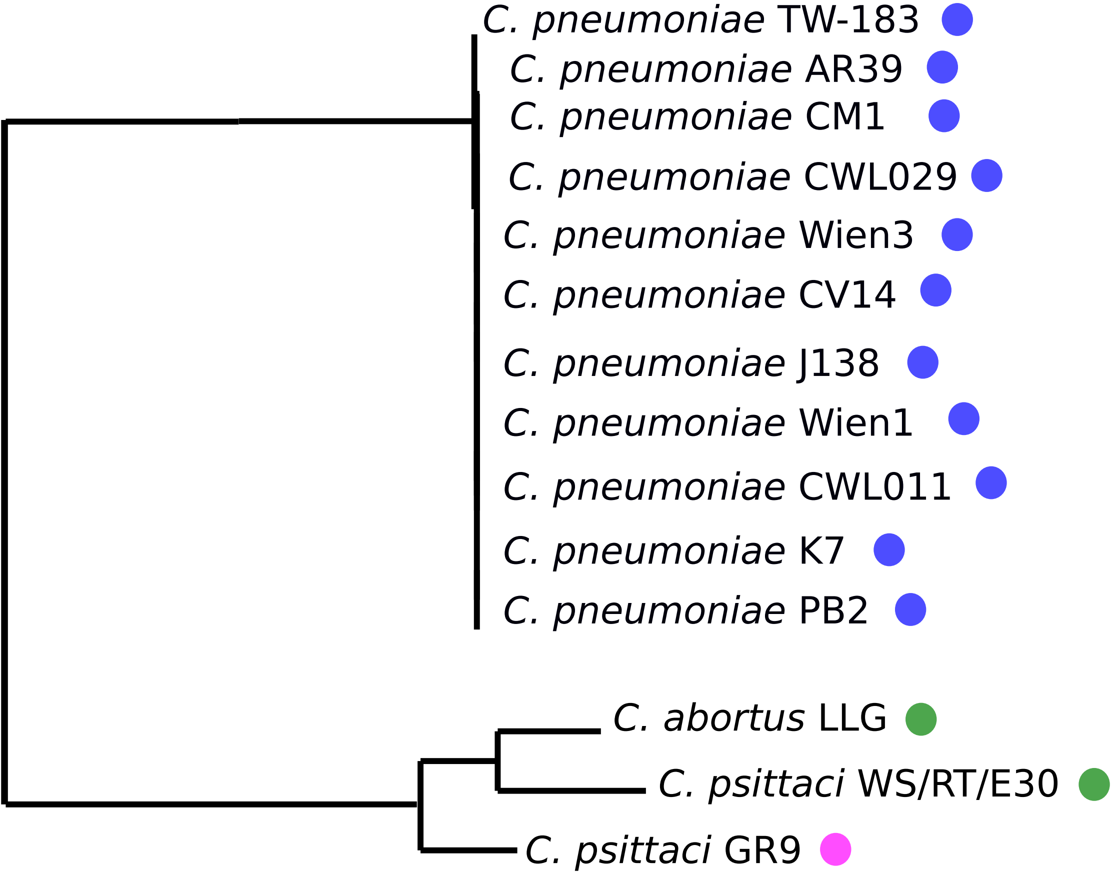

### Supplemental file Figure 4

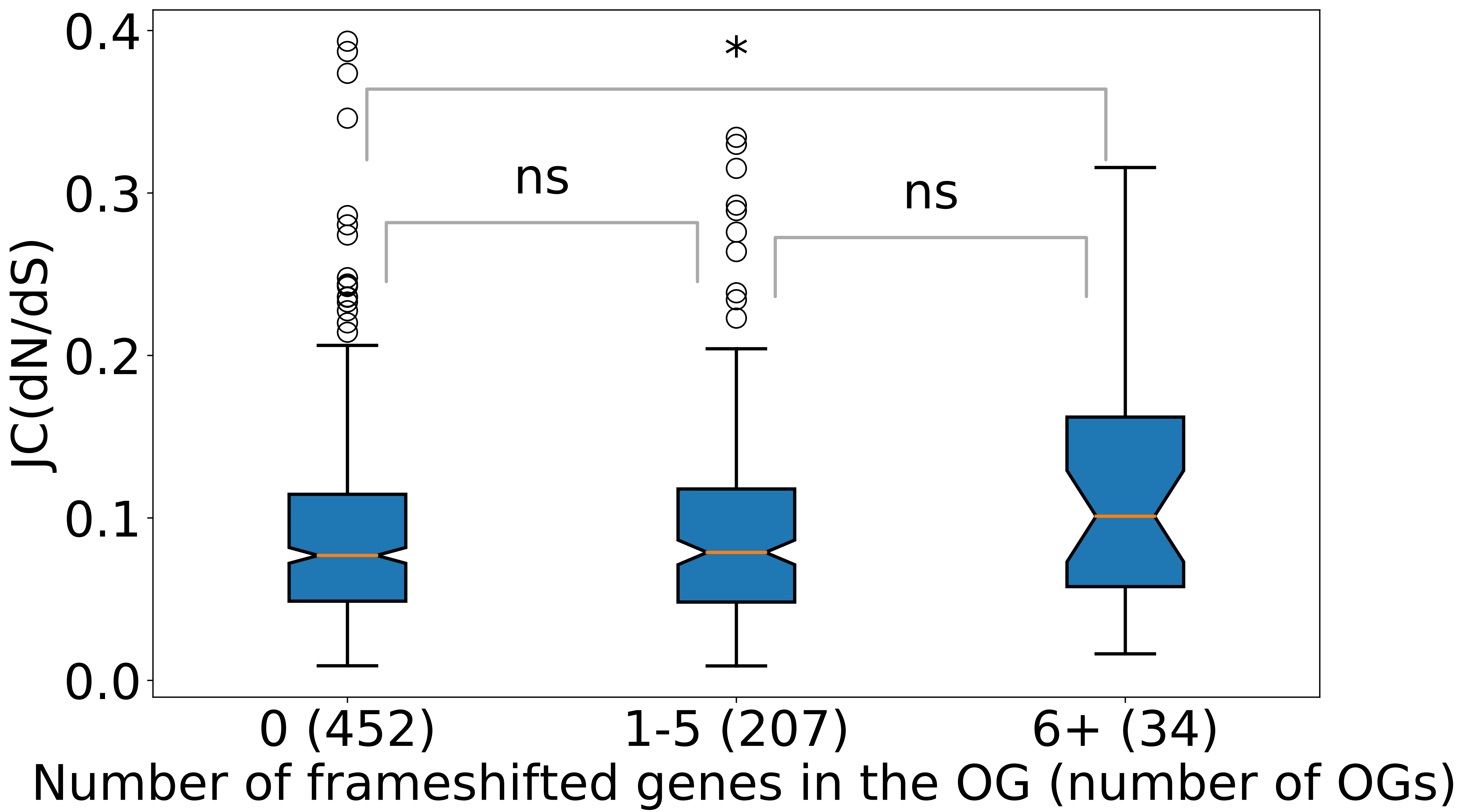

### Supplemental file Figure 5

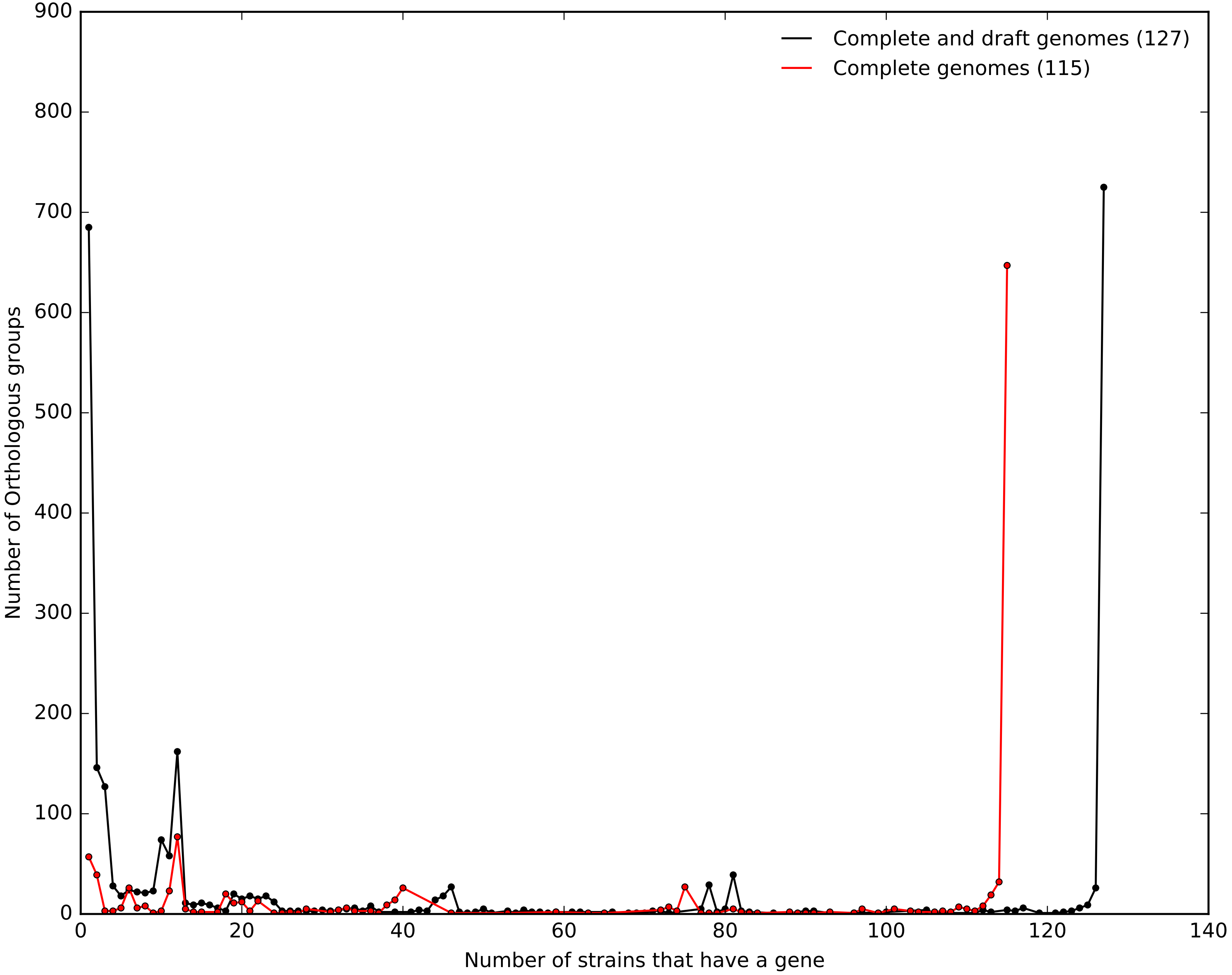

### Supplemental file Figure 6

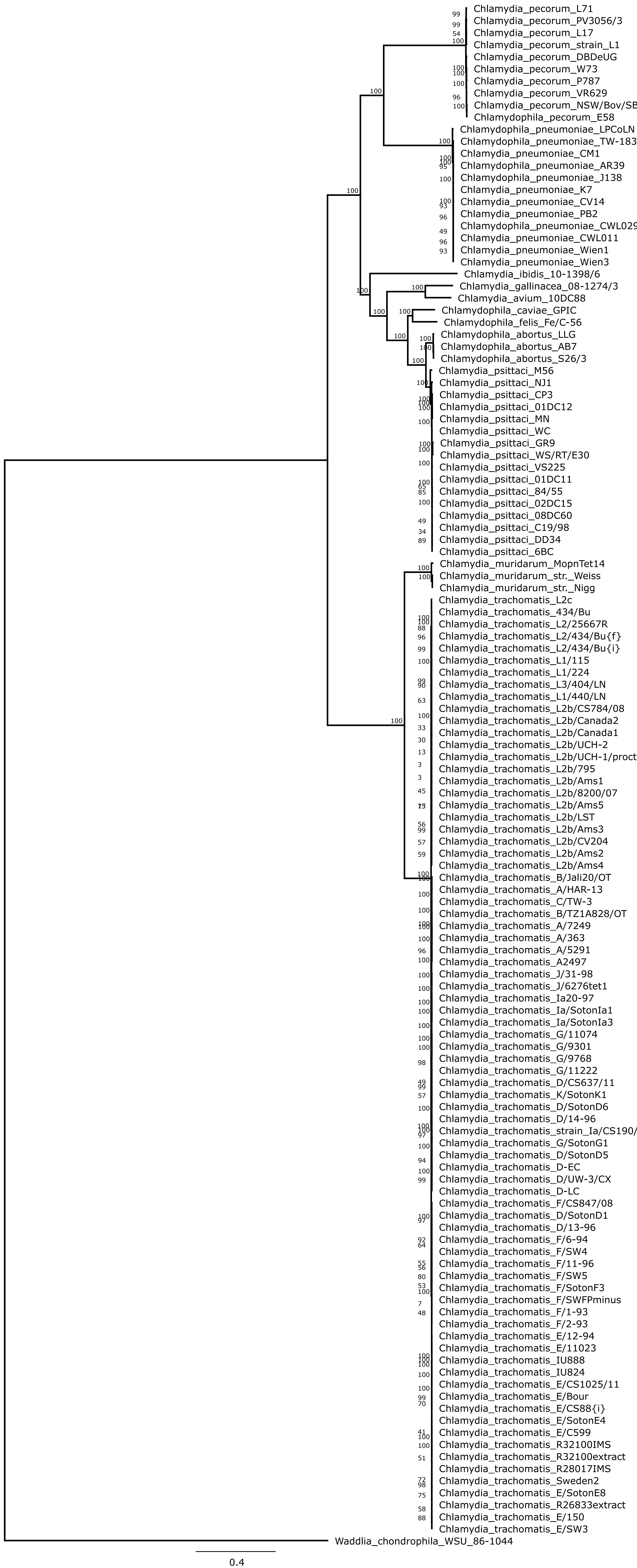
